# Simulating future rewards: exploring the impacts of implicit context association and arithmetic booster in delay discounting

**DOI:** 10.1101/2020.03.03.974709

**Authors:** Minho Hwang, Sung-Phil Kim, Dongil Chung

## Abstract

People have higher preference for immediate over delayed rewards, and it is suggested that such an impulsive tendency is governed by one’s ability to simulate future rewards. Consistent with this view, recent studies have shown that enforcing individuals to focus on episodic future thoughts reduces their impulsivity. Inspired by these successful reports, we hypothesized that administration of a simple cognitive task which is linked with future thinking might also function as an effective tool for modulating individuals’ preference for immediate (or delayed) rewards. Specifically, we used one associative memory task and one working memory task that each of which was administered to intervene acquired amount of information and individuals’ ability to construct a coherent future event, respectively. Among the set of cognitive tasks, we found that only the arithmetic working memory task had a significant effect of reducing individuals’ impulsivity. However, in our follow-up experiment, this result was not replicated. Across these two independent experiments, we observed a significant main effect of repetition in individuals’ impulsivity measure, such that participants showed more patient choices at the second compared to the first assessment task. In conclusion, there was no clear evidence supporting that our suggested intervention tasks effectively reduce individuals’ impulsivity, while the current results call attention to the importance of taking into account task repetition effects in studying the impacts of cognitive training and intervention.

## 1. Introduction

In daily decision-making, people sometimes make a choice not because it is better now, but because it will bring them a higher return in the future. One’s exaggerated preference for the temporally proximate small reward against the delayed large reward has been referred to as impulsivity (Ainslie, 1975; Baker, Johnson, & Bickel, 2003; Bickel & Marsch, 2001; Kable & Glimcher, 2007b), and the choices that show the opposite pattern—decisions with consideration of future consequences—are considered as outcomes of successful self-control (Figner et al., 2010; Hare, Camerer, & Rangel, 2009; McClure, Laibson, Loewenstein, & Cohen, 2004). A large body of studies examined the tradeoffs between the time of reward delivery and the amount of reward, and suggested that discounting the value of delayed rewards explains why typical individuals often show seemingly impulsive behaviors, i.e., to choose the small immediate reward over the large delayed reward (Frederick, Loewenstein, & O’donoghue, 2002; Green & Myerson, 2004; Mazur, 1987). This framework of temporal discounting in valuation extends to real-life health risk behaviors, such as substance abuse (Baker et al., 2003; Bickel & Marsch, 2001; MacKillop et al., 2011) and problem gambling (Dixon, Marley, & Jacobs, 2003; Petry & Casarella, 1999). Consistent with the view that one discounts delayed rewards, individuals who tend to make choices that may deliver immediate pleasure than longterm healthiness (e.g., eating a chocolate bar than an apple as dessert) also showed higher discounting rates in decision-making tasks (Bickel et al., 2014; Story, Vlaev, Seymour, Darzi, & Dolan, 2014).

Why do some individuals discount delayed rewards more than others in valuation process? Recent studies found evidence suggesting that one’s ability to simulate future rewards is linked with the extent to which one is sensitive to the temporal delay of rewards (Benoit, Gilbert, & Burgess, 2011; Hakimi & Hare, 2015; Peters & Büchel, 2010; Schacter et al., 2012); individuals’ preference for delayed large rewards significantly increased when they were explicitly instructed to imagine the reception of the reward in association with a cue selected from their episodic future plans (e.g., trip to Paris). These findings supported that episodic future thinking induces vivid imagination (simulation) of future reward delivery, and in turn, reduces reward delay discounting. However, the mechanisms via which the induction successfully reduces individuals’ delay discounting remains unknown.

Here, we focus on two among numerous potential factors that are closely related to future reward simulation and thus may be enhanced in the process of episodic future thinking (D’Argembeau, Ortoleva, Jumentier, & Van der Linden, 2010; Schacter et al., 2012): a particular association between reward information with a context and individuals’ working memory ability. Specifically, the following two hypotheses were tested. First, additional context information (e.g., specific type of event) associated with the delayed reward will increase certainty of the simulated contents and may help individuals simulate the reward’s subjective value (Szpunar, 2010). Second, one’s working memory capacity may be crucial in simulating (estimating) the reward’s subjective value across temporal horizon, such that increased working memory capacity would facilitate the simulation process (Hill & Emery, 2013; Zavagnin, De Beni, Borella, & Carretti, 2016). These hypotheses were tested by examining whether modulation of each factor indeed significantly reduced individuals’ delay discounting, which is an indication of better simulation of delayed rewards. In our first experiment, our results provide evidence with medium to large effect size that enhancement in one’s working memory capacity is associated with reduction of her delay discounting. However, in an independent second experiment, we report that this intervention effect is not replicable. Finally, across the two experiments and regardless of the type of cognitive modulation that was applied, in contrast to our expectation, our data show a significant main effect of repetition in individuals’ impulsivity measure.

To test the impacts of modulating each factor on reducing individuals’ delay discounting, we first administered a classic intertemporal choice task (ITC task; adapted from Kable & Glimcher (2007a)) (**Fig. 1a, S1a**) and measured individuals’ baseline delay discounting. During the ITC task, participants made a series of choices between a fixed immediate reward of 10,000 KRW (around $10) and a larger delayed reward that varied trial to trial (see **4.2. Experimental procedures** for detailed experimental procedures). Based on participants’ choices across three sessions of the ITC task, we estimated individuals’ delay discount rate (*k*); we assumed that an objective reward V delivered in a delay of D is subjectively valued following a hyperbolic discount function SV = V/[1 + *k*D] (Azfar, 1999; Frederick et al., 2002; Green & Myerson, 2004; Laibson, 1997).

**Figure 1.**
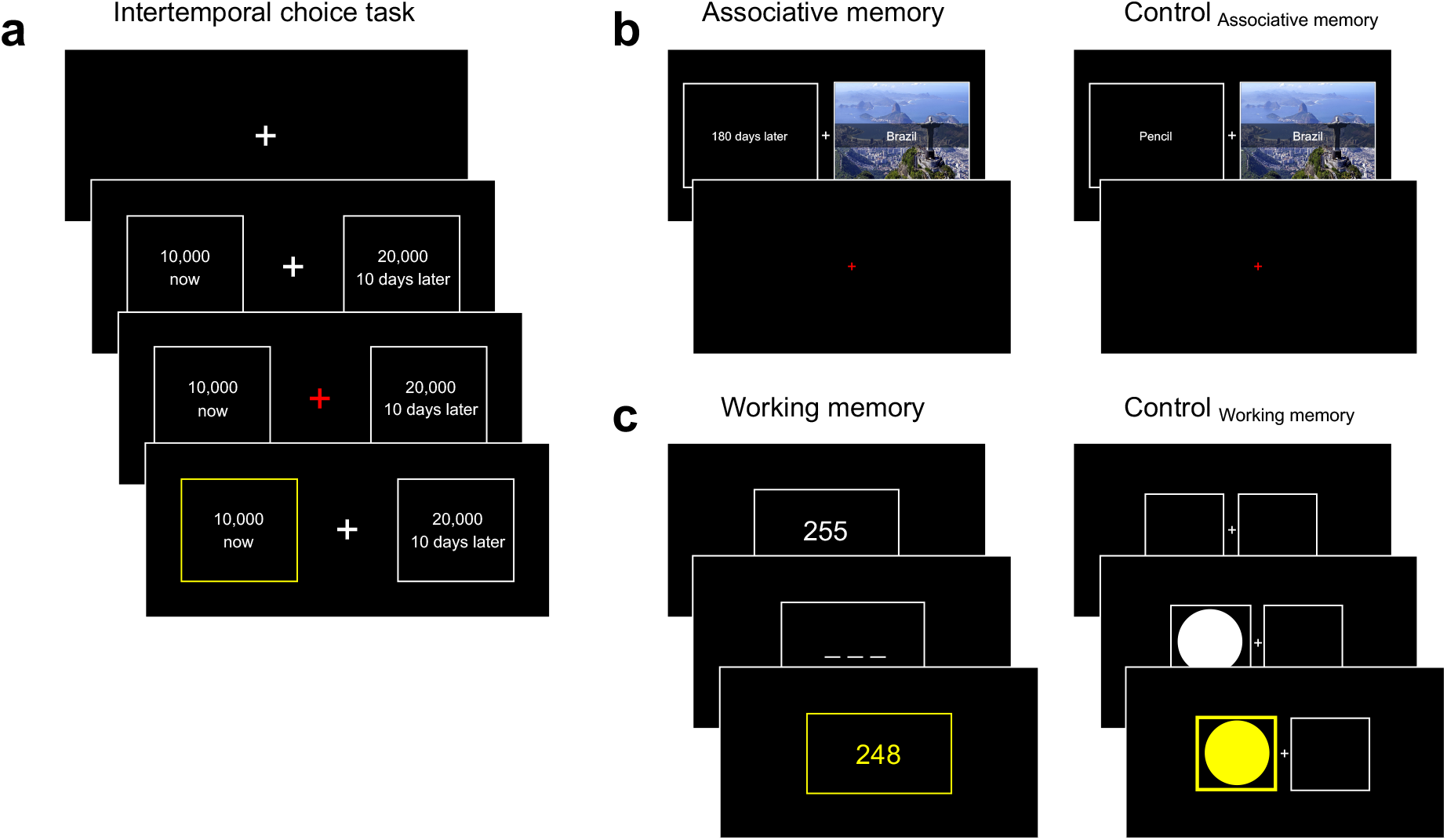
The intertemporal choice task and cognitive intervention paradigms. **(a)** During the intertemporal choice (ITC) task, participants made choices between an immediate, smaller reward and a delayed, larger reward. The immediate reward was fixed (10,000 Korean Won (KRW)), but the alternative delayed option was constructed using six distinct delays (1, 10, 21, 50, 90, and 180 days) and varying size of delayed rewards (10,300-29,300 KRW) determined by stepwise approach. Individuals could make a response when the crosshair turned red, and a feedback screen followed their choices to highlight the chosen option. **(b, c)** Two intervention tasks and their control tasks were used to examine impacts of associative and working memory on future reward simulation, and delay discounting in turn. **(b, left)** Associative memory task comprised encoding and retrieval phases (not depicted here). In the encoding phase, participants were asked to memorize associations between the paired stimuli. Specifically, names of the countries with a picture of their representative must-see sights were matched with ‘expected preparation time’ to go on a trip to the places. Note that the preparation time was presented in the same format as the delays in the ITC task. **(b, right)** Control_AM_ was largely identical to the Associative memory task, except that the associations were made between country names and various stationaries. **(c, left)** In Working memory task, participants had to remember a random three digit number presented on the screen (e.g., 255), mentally subtract seven, and report the number on the subsequent screen (e.g., 248). They were asked to repeat this mental calculation (e.g., 241, 234, 237, …) until a new random number was provided. **(c, right)** During Control_WM_, participants had to maintain sustained level of attention. Specifically, on each trial, a circle showed up on either left or right half of the screen, and participants were asked to press the arrow key that matches the location as soon as possible.

To examine whether individuals’ impulsivity can be modulated, we introduced one of five types of intervention tasks (Associative memory task, Working memory task, and three control tasks) after the baseline ITC task, and then a second assessment of ITC task (**Experiment 1**). To examine whether the results from Experiment 1 are replicable, we conducted another independent experiment where we tested three specific types of intervention tasks among the original five (**Experiment 2**).

## 2. General methods

### 2.1. Participants

216 healthy young adults (male/female = 124/92) participated in the current study. None of the participants reported a history of neurological or psychiatric illness. Two independent studies were conducted and there were no overlapping participants across two experiments. 126 healthy individuals (male/female = 71/55, age = 22.00±2.48) participated in Experiment 1, conducted between March 25-May 20, 2019, and 90 individuals (male/female = 53/37, age = 22.24±3.93) were recruited for Experiment 2, conducted between January 13-July 19, 2021. All participants provided written informed consent and were paid for their participation. The study was approved by the Institutional Review Boards of Ulsan National Institute of Science and Technology (UNISTIRB-18-18-A).

### 2.2. Experimental procedures

The aim the current study was to measure individuals’ impulsivity, and examine whether intervention of cognitive factors alter their characteristics. At the beginning of each individual’s participation, he or she was randomly assigned to one of the five subgroups (three subgroups for Experiment 2), each of which was paired with one type of intervention task. All participants completed one session of Intertemporal choice task (ITC), one session of the assigned intervention task, and one additional session of ITC. Participants were paid at the end of the study, based on the delay and reward of a random single option drawn from all the ITC task decisions the participant made. All experiments were performed in accordance with relevant guidelines and regulations.

#### 2.2.1. Intertemporal choice task

During the ITC task, participants made a series of choices between a fixed immediate reward of 10,000 KRW (around $10) and a larger delayed reward that varied trial to trial (**Fig. 1a, S1a**). Six distinct delays (1, 10, 21, 50, 90, and 180 days) were used for the delayed reward option, and the value of the option was determined following the staircase approach. There were two unique initial values for the large delayed reward (15,000 and 20,000 KRW) and the staircase approach is taken for each pair of [delay, initial value]. Specifically, if a participant chose the immediate option at n^th^ step, reward of the delayed option was set to [delayed reward at n^th^ step – 10,000 × (1/2)^n-1^] (n = 1, 2, 3, 4, 5) for the next trial where the same pair of [delay, initial value] was used. On the contrary, if a participant chose the delayed option, reward of the delayed option was set to [delayed reward at n^th^ step + 10,000 × (1/2)^n-1^] for the next trial where the same pair of [delay, initial value] was used. There was one exception. To avoid the delayed reward being equal to or smaller than the immediate reward (10,000 KRW), choices of the delayed option on the trials under the initial value of 15,000 KRW led to the next trial with delayed reward of [delayed reward at n^th^ step – (delayed reward at n^th^ step – 10,000) × (1/2)^n-1^]. This ‘titration’ procedure repeated for five iterations (steps) for each distinct pair of delay and initial value. Each block comprised 60 trials ([6 delays] × [2 unique initial values for the large reward (15,000 and 20,000 KRW)] × [5 iteration staircase]). To minimize the effect of participants remembering previous pairs of options and their choices, the trial order was randomized for delay × initial value with a unique order per participant. Because of the staircase approach, the subsequent trial was randomly chosen among 12 available pairs [6 delays × 2 initial values] including the pair that was present on the current trial. Again, the amount of delayed reward was determined by the aforementioned rule. Participants completed three blocks of ITC task, 180 trial in total ([6 delays × 2 initial values × 5 steps] × 3 blocks).

#### 2.2.2. Intervention tasks

Previous studies suggested that episodic future thinking induces vivid imagination (simulation) of future reward delivery, and in turn, reduces reward delay discounting (Benoit et al., 2011; Hakimi & Hare, 2015; Peters & Büchel, 2010; Schacter et al., 2012). To investigate potential mechanisms underlying this induction effect, we examined impacts of two factors that are known to be linked with episodic future thinking (or mental simulation): acquisition of additional context information and enhancement of individual working memory performance. Two intervention tasks were designed and used to examine these factors, along with three intervention tasks as controls. In the next section, we explain detailed designs of each intervention task.

#### 2.2.3. Intervention task: Associative memory task

During Associative memory task, participants were asked to memorize the association between temporal delays and names of the countries, which were introduced as pairs of information about the most wanted places to visit and average duration people spend planning for the trip (**Fig. 1b, S1bi**). This intervention was to implicitly provide additional information about delayed reward options; if additional context information benefits simulating delayed rewards, the intervention should decrease individuals’ delay discounting in the subsequent ITC task.

Associative memory task comprised two phases: encoding and retrieval phases. Name of 30 different countries and 30 unique temporal delays (preparation time for the trip) were used as pairs. Among the pairs, only six levels of delays were shared with the delays in ITC (1, 10, 21, 50, 90, and 180 days) and other 24 levels of delays were never used in ITC. The name of the countries were sorted by their geographical distances from South Korea in ascending order, and they were matched with delays accordingly; examples include [Japan, 1 day], [Vietnam, 10 days], [Taiwan, 21 days], [Philippine, 50 days], [New Zealand, 90 days], and [Brazil, 180 days]. During the encoding phase, there were 60 trials where 30 pairs that should be memorized were presented twice. On each trial, after presentation of the association, participants had to report whether they could successfully remember and vividly image the association between the paired pieces of information.

During the retrieval phase, participants were asked to retrieve previously memorized association pairs and verbally report answers. Specifically, each retrieval trial was cued by one of the 30 delays, and participants’ verbal answers of the associated country names were recorded for the accuracy assessment. On each trial, participants used a keypress to notify that they finished providing the answer, and then they reported the extent to which they were confident about the answer in a 5-likert scale (1 = not confident and 5 = very confident). The target delays were tested three times, while the non-target delays were tested only once. In total, there were 42 trials where the target and non-target delay cues were intermixed.

#### 2.2.4. Intervention task: Control for Associative memory task (Control_AM_)

The procedures of the Control_AM_ task were largely identical to those of the Associative memory task (**Fig. 1b, S1bii**). The only difference was that various office supplies items were paired with the country names in replacement of delays. Moreover, to match cognitive workload with the Associative memory task, office supplies items were used that word length of which names (in Korean) matched the number of digits of the delay it is replacing. We chose office supplies items instead of delays, so that no cues during the ITC may trigger retrieval of the information regarding the learned association.

#### 2.2.5. Intervention task: Working memory task

During Working memory task, participants were asked to conduct a series of mental calculation (**Fig. 1c, S1ci**). Specifically, participants played a task so called ‘Serial 7s subtraction (Hayman, 1942)’ where they had to subtract seven from the answer of the previous trial. For example, if the answer of the previous trial was 200, the next answer that should be entered is 193. At the beginning, an initial number was randomly chosen between 107 and 999, and presented on the screen. Participants had to remember the most recently entered answer, so that they could use the information for the subsequent trial. Such needs for reservation of information for a short duration is known to recruit participants’ working memory (Klingberg, 2010). There were three conditions where the task reset with a new initial number: when participants entered wrong answer, when the answer for the subsequent trial was smaller than 100, or when participants did not enter an answer within 20 seconds. Participants completed two blocks of task where each block lasted for 10 minutes.

#### 2.2.6. Intervention task: Control for Working memory task (Control_WM_)

Control_WM_ task was designed to require continuous attention from participants, but not working memory (**Fig. 1c, S1cii**). On each trial, a circle shaped stimulus showed up on either left or right half of the screen, and participants were instructed to press a left or right arrow key that matches the side of the screen. Stimulus color changed to yellow immediately after a keypress only if the keypress were correct, the color changed to gray when no answers were entered within 1.5 seconds, but no change was made when the submitted answer was incorrect. The next trial started after a short feedback screen (0.5 seconds) and a jittered inter-trial interval (U(0.3, 0.7)). Participants completed two blocks of task where each block lasted for 10 minutes.

#### 2.2.7. Intervention task: Resting

To examine the impacts of intervention tasks, all participants went through two sessions of ITC tasks, one before and one after the corresponding intervention. To control for a possibility that the impulsivity change may be induced by the repeated participation in ITC task, we introduced a resting phase for one of the five subgroups. The length of the resting phase was set to 20 minutes, matching the average length of other tasks.

### 2.3. Behavioral analyses

For each subgroup, participants’ tendency of choosing the delayed option over the immediate option was calculated across three blocks of the ITC task, separately for the task before and after a intervention task. As a model-agnostic measure, the impact of each intervention was calculated as a difference of the choice ratio between the two choice ratio measures. In addition, we estimated individuals’ delay discounting rate (*k*) based on participants’ choices across three blocks of the ITC task, separately for the task before and after an intervention. We assumed that an objective reward V delivered in a delay of D is subjectively valued following a hyperbolic discount function SV = V/[1 + *k*D], and that softmax choice rule explains the link between subjective valuation and choices as below:

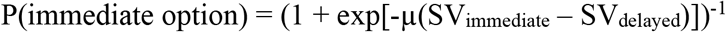

where SV_immediate_ (SV_delayed_) is the subjective value of the immediate (delayed) reward option and μ is the inverse temperature (or sensitivity to value differences). Individual parameter estimations were conducted with custom MATLAB R2017a scripts using maximum loglikelihood estimation (MLE) at the individual subject level. The *fminsearch* function in MATLAB with different initial values starting estimation was used. After the parameter estimation, changes of delay discounting rates in each subgroup were used as model-based measures of the impact of the intervention tasks.

To examine whether the extent to which individuals successfully completed their assigned intervention task was associated with the change of behavioral choice patterns in the ITC task, performance of intervention tasks was measured accordingly. First, mean accuracies were calculated for each task. Second, for the Working memory task and Control_WM_, temporal evolution in individuals’ performance was also examined. To do this, these tasks were binned into equal size (10 trials per bin) using sliding window (5-trial overlap). We examined whether participants showed increasing performance across bins, which would be an indication of successful training of the corresponding cognitive function. Furthermore, to better illustrate the impacts of intervention tasks on the changes in individuals’ impulsivity (i.e., delay discounting rate), we divided each intervention task in five larger bins (each bin’s size was 1/3 of the entire task length where sliding window was used with overlapping 1/6 of task length) and the association within each bin was calculated.

### 2.4. Statistical analyses

The size of each intervention task group was relatively small, and thus any descriptive measures may be sensitive to a particular outlier. To prevent such biases, we used a bootstrapping method with 10000 resampling iterations for within- and between-group mean comparisons. P-value for each statistical analysis was calculated as the proportion of extreme samples (based on t-values) out of the entire resample estimates, relative to the statistics estimated from the original data set. The same bootstrapping approach was used for regression analyses between intervention task performances and ITC behavioral patterns.

## 3. Experiment 1

### 3.1. Methods

In Experiment 1, 126 healthy individuals (male/female = 71/55, age = 22.00±2.48) were recruited and were randomly assigned to one of the five intervention tasks: two hypothesis-driven tasks—i) association memory task and ii) working memory task—and three control tasks (see **2.2. Experimental procedures** for details). The Associative memory task was designed to implicitly provide additional context information about delayed rewards (**Fig. 1b, S1b**), and the Working memory task was to boost individuals’ working memory capacity (**Fig. 1c, S1c**). Specifically, by using the Associative memory task intervention, we examined the possibility that the memorized association would be retrieved during the ITC task due to the use of common cues (i.e., delays for reward delivery vs. delays for planned trips; see **2.2. Experimental procedures** for details), and that this additional information (i.e., planned trip locations) would provide contexts facilitating the simulation of future rewards (Peters & Büchel, 2010). By using the Working memory task, we tested the hypothesis whether or not increase in working memory capacity would reduce one’s delay discounting. This hypothesis was constructed based on previous reports suggesting that mental calculation shares the same cognitive resources with time perception (Brown, 1997), which provides a crucial piece of information in future planning. Moreover, it is known that working memory serves to integrate information across multiple dimensions in mental simulation (Hill & Emery, 2013; Szpunar, 2010). Thus, we expect that enhancing one’s working memory capacity may facilitate one’s future reward simulation.

Three control tasks comprised i) another association task (Control_AM_) where participants learned association about information irrelevant to delays (**Fig. 1b, S1b**), ii) a spatial attention task (Control_WM_) where participants were required of using continuous attention, but not working memory (**Fig. 1c, S1c**), and iii) resting (see **2.2. Experimental procedures** for task details). A second ITC task was administered to all participants after each type of intervention, and the discount rate changes were calculated to examine the impacts of modulating each factor on reducing individuals’ delay discounting. Participants were paid at the end of the study based on the delay and reward of a random single option drawn from all the ITC task decisions the participant made.

26 students participated in the Associative memory task, and 24 students participated in the Working memory task. For control tasks, comparable number of students were assigned who were age, sex, and education matched (Control for Associative memory task (Control_AM_): N = 25; Control for Working memory task (Control_WM_): N = 24; Resting: N = 26). From the entire participant pool, we excluded six participants from analyses because they explicitly expressed that they would rather like to receive class credits than monetary rewards as a compensation for participation; we considered this as an indication of different, subjective incentive structure, and thus excluded these participants. One additional participant was excluded from the analyses because he showed up at the lab under the influence (drunk). Additional eight participants were excluded for whom individual-level computational model (see **2.3. Behavioral analyses**) did not arrive at a unique solution using maximum likelihood estimation (MLE). After exclusion, data from 111 participants (male/female = 60/51, age = 22.07±2.54) were used for analyses (see **Table 1** for demographic information for each subgroup).

**Table 1.** Demographic data for Experiment 1.

|  | Associative<br>memory<br>(N = 23) | Control<sub>AM</sub><br>(N = 22) | Working<br>memory<br>(N = 22) | Control<sub>WM</sub><br>(N = 21) | Resting<br>(N = 23) |
| --- | --- | --- | --- | --- | --- |
| Age (years)<sup>a</sup> | 22.22±2.71 | 21.45±2.15 | 22.32±2.44 | 22.57±2.96 | 21.83±2.46 |
| Male/female<sup>b</sup> | 11/12 | 12/10 | 17/5 | 11/10 | 9/14 |
| Education level<sup>c</sup> |  |  |  |  |  |
| Some college | 17 | 19 | 19 | 15 | 18 |
| Finished college | 6 | 3 | 3 | 6 | 5 |

Five subgroups were of comparable age<sup>a</sup> ( $F(4, 106) = 0.64, P = 0.63$ ), gender<sup>b</sup> ( $\chi^2(4) = 5.08, P = 0.28$ ), and education<sup>c</sup> ( $\chi^2(4) = 2.55, P = 0.64$ ).

### 3.2. Results

All participants completed the first assessment of ITC task to have their individual baseline discount rates estimated (log transformed *k*; see **2.3. Behavioral analyses** for parameter estimation procedures). At this first assessment, the five sub-groups showed comparable baseline delay discount rates (F(4, 105) = 0.43, *P* = 0.78, one-way ANOVA; **Fig. 2**). This allowed us to define the impact of each intervention as the behavioral differences between pre- and post-intervention tasks; we calculated Δlog *k* = log(*k*_pre-intervention_) – log(*k*_post-intervention_) for model-based analyses where *k* indicates an individual’s estimated discount rate (larger *k* refers to more impulsive behavioral choices). In the measure Δlog *k*, positive numbers indicate a positive intervention effect on enhancing preference for the delayed reward option (i.e., reducing impulsivity).

**Figure 2.**
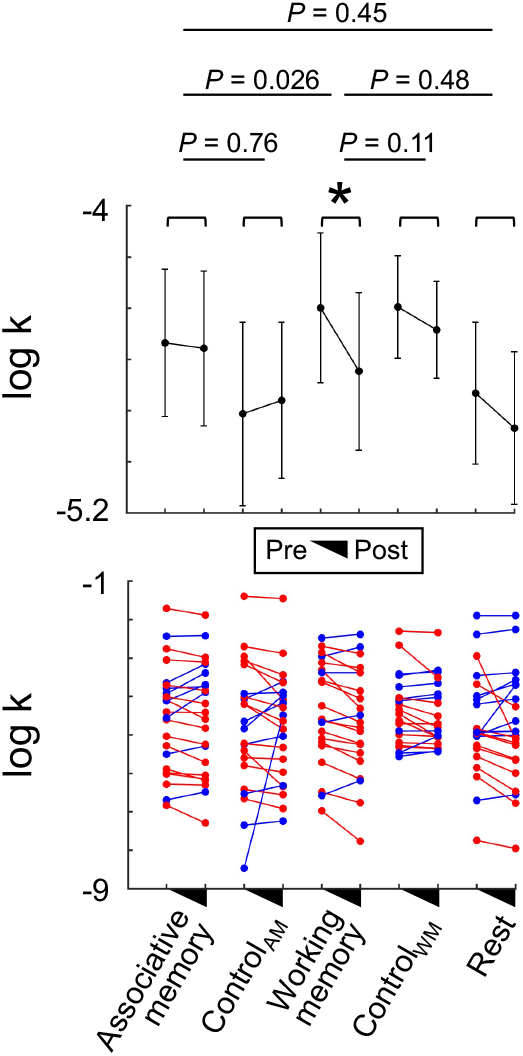
Impacts of the intervention tasks on reducing individuals’ impulsivity (Experiment 1). Each participant group was given with a different type of intervention task. To examine whether each intervention had an impact on individuals’ impulsivity, we calculated changes in individuals’ delay discount rates (*k*) by comparing their behavioral patterns during the intertemporal choice (ITC) task conducted before and after the intervention task. Individuals showed significantly reduced impulsivity after participating in the Working memory task (Paired t-test, t(20) = 3.44, *P* = 0.0026, Cohen’s d = 0.75, bootstrapping *P* = 0.0031). On the contrary, individuals who participated in other intervention tasks did not show the same pattern (i.e., reduced impulsivity) (all bootstrapping *p*s>0.05). The mean discount rate change in individuals who participated in the Working memory task was larger compared with that in individuals who were assigned to Associative memory task in between two ITC tasks (Independent-sample t-test, t(42) = −2.31, *P* = 0.026, Cohen’s d = −0.35, bootstrapping *P* = 0.026). The impact of the Working memory task was not statistically different from that of the corresponding control task (Control_WM_; bootstrapping *P* = 0.11) nor the resting (bootstrapping *P* = 0.48). Error bars indicate s.e.m.; \**P* < 0.05.

As explained above, choices from individuals who were assigned to the Associative memory task and to the Working memory task were used to test the impacts of the context information and the working memory capacity on reducing impulsivity, respectively (**Fig. 2**). A mixed-design ANOVA setting ‘Time’ (pre- and post-intervention) as a within-group factor and ‘Type’ (Associative memory, Working memory, and Rest) as a between-group factor revealed a significant main effect of Time (F(1, 64) = 5.79, *P* = 0.019). However, the main effect of Type and the interaction of Time × Type were not significant (Type: F(2, 64) = 0.28, *P* = 0.75; Interaction: F(2, 64) = 1.39, *P* = 0.26), suggesting that neither intervention task had superior efficiency in reducing individuals’ impulsivity compared to simple repetition of ITC (i.e., the Rest group). The absence of significant differences among the impact of tasks could be due to insufficient statistical power. Alternatively, there is a possibility that the impact of each intervention may be tightly intertwined with their performances at the cognitive intervention task. We further examined this possibility for the sake of the purpose of the current study to explore the impacts of potential intervention mechanisms linked with simulating future rewards.

Within each intervention group, Working memory task was the only intervention that had a significant impact between two ITC measures, such that individuals showed significantly reduced temporal discounting rate after the intervention (Paired t-test, t(20) = 3.44, *P* = 0.0026, Cohen’s d = 0.75, bootstrapping *P* = 0.0031; **Fig. 2**). Individuals who took part in Associative memory task as an intervention and who rested between two measures of ITC all did not show difference in their estimated temporal discounting rate (Associative memory: t(22) = 0.34, *P* = 0.74, Cohen’s d = 0.070, bootstrapping *P* = 0.74; Rest: t(22) = 1.049, *P* = 0.31, Cohen’s d = 0.22, bootstrapping *P* = 0.31; **Fig. 2**). Note that neither of the two additional tasks (Control_WM_ and Control_AM_), each of which was examined as a control for the Working memory and Associative memory interventions, respectively, were effective in reducing individuals’ delay discounting rates (Control_WM_: t(20) = 1.40, *P* = 0.18, Cohen’s d = 0.31, bootstrapping *P* = 0.19; Control_AM_: t(21) = −0.25, *P* = 0.80, Cohen’s d = −0.054, bootstrapping *P* = 0.81).

Direct comparisons between the intervention groups showed that the reduction of delay discounting rate in the Working memory group was significantly larger than that in the Associative memory group (Two-sample t-test, t(42) = 2.31, *P* = 0.026, Cohen’s d = 0.35, bootstrapping *P* = 0.026; **Fig. 2**). Although trending toward the superior impact, the impact of the Working memory intervention was comparable with that of the Control_WM_ (t(40) = 1.65, *P* = 0.11, Cohen’s d = 0.25, bootstrapping *P* = 0.11) and that of the Rest (t(42) = 0.73, *P* = 0.47, Cohen’s d = 0.11, bootstrapping *P* = 0.48; **Fig. 2**). The impact of the Associative memory intervention was also comparable with that of its corresponding control tasks (Associative memory vs Control_AM_: t(43) = 0.35, *P* = 0.73, Cohen’s d = 0.052, bootstrapping *P* = 0.76; vs Rest: t(44) = −0.78, *P* = 0.44, Cohen’s d = −0.12, bootstrapping *P* = 0.45; **Fig. 2**). These results all together suggest that individuals’ working memory capacity, but not abundancy of context information or individuals’ attention level, is linked with their preference for delayed rewards. Yet again, given the statistically comparable impacts of the intervention tasks, one should be careful not to overinterpret the involvement of working memory in simulation of future rewards.

For completeness, we also compared the impacts of each intervention design using model-agnostic measures; ΔP(immediate option) = P(immediate option_pre-intervention_) – P(immediate option_post-intervention_) was calculated as a model-agnostic measure where P(immediate option) refers to probability of choosing the immediate option. Note that the model-based measure, delay discounting rate *k*, is more appropriate in quantifying individuals’ characteristics of future reward simulation, because the model-agnostic measure cannot tease apart the character-of-interest (i.e., delay discounting rate) from a confounding factor (e.g., value sensitivity, denoted as μ; see **2.3. Behavioral analyses**). Nevertheless, the statistical results were largely the same as model-based results (see **Fig. S2** for statistical results).

We further examined whether or not individuals’ performance in the Working memory task is associated with the level of impulsivity reduction. Repetition of working memory tasks typically results in improvement of individuals’ task performance, which is considered as a signature of cognitive training (Bickel, Yi, Landes, Hill, & Baxter, 2011; Sarah E Snider et al., 2018). Although our Working memory task as a type of intervention task was only in the order of tens of minutes, rather than days as in other cognitive training studies, we also observed such a performance enhancement across time (**Fig. 3a**). Specifically, 91% of participants showed performance increase along the task and across individuals, proportion of correct answers significantly increased along the course of the task (r = 0.64, *P* = 2.97e-5). To examine the impact of this intervention task on individuals’ impulsivity, we examined whether or not the average accuracy and/or the speed of performance improvement were associated with the change of impulsivity rate.

**Figure 3.**
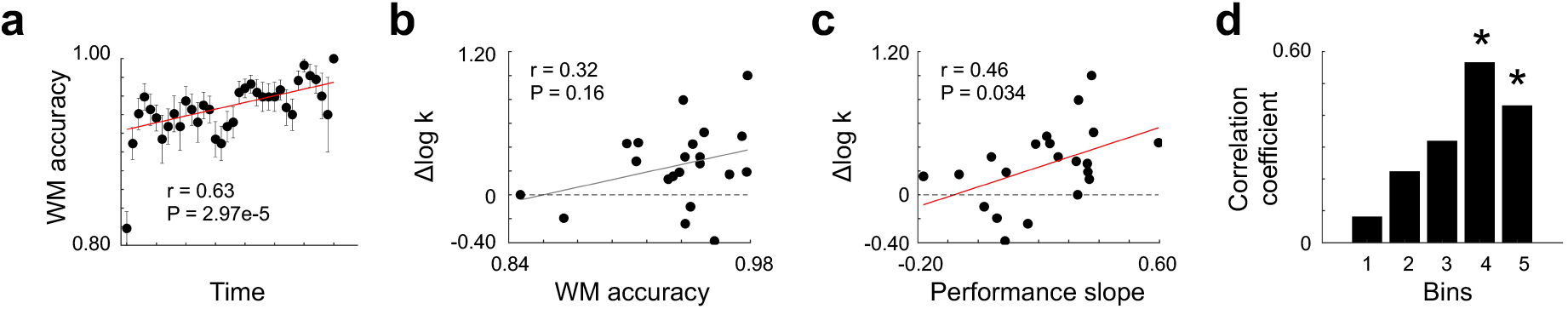
Individual performance enhancement in working memory training is associated with the level of impulsivity reduction (Experiment 1). **(a)** Change over time in Working memory task accuracy was measured to examine whether or not individuals showed performance enhancement. Average accuracy within each 10-trial sliding window (with 5-trial overlap) was calculated per participant. Indeed, participants showed a significant performance improvement over time (Pearson’s correlation r = 0.64, *P* = 2.97e-5). Each dot represents mean accuracy within each window across participants, and error bars indicate s.e.m. **(b)** Although statistically not significant, individuals’ overall performance (i.e., accuracy) in Working memory task had a trend of positive correlation with the changes in their delay discounting rates (r = 0.32, *P* = 0.16, bootstrapping *P* = 0.16). Specifically, individuals who had the highest (lowest) accuracy in Working memory showed relatively larger (smaller) reduction in their estimated impulsivity (positive Δlog *k* indicates one’s impulsivity reduction). **(c)** Independent of individuals’ initial accuracy level, individuals’ speed of Working memory performance enhancement (performance slope) was significantly correlated with the level of impulsivity reduction (r = 0.46, *P* = 0.036, bootstrapping *P* = 0.034). Each dot represents an individual, and solid red lines are robust regression lines. **(d)** By dividing the Working memory task into five bins (bin size = 1/3 of the entire task length; sliding window with 1/6 overlap), a stark difference was observed between bins in the association between individuals’ Working memory accuracy and their impulsivity reduction levels. Particularly, individuals’ Working memory task performance in the later period (fourth and fifth bins) showed strong associations with the extent to which they make less impulsive choices in the post-intervention ITC task. *\*P* < 0.05.

There was a positive, but statistically non-significant trend between average working memory performance and reduction of delay discounting (Pearson correlation, r = 0.32, *P* = 0.16, bootstrapping *P* = 0.16; **Fig. 3b**). On the contrary, participants who had steeper improvement of the working memory performance showed significantly larger reduction in their impulsivity (r = 0.46, *P* = 0.036, bootstrapping *P* = 0.034; **Fig. 3c**). This result was corroborated with the association between impulsivity reduction and average working memory performance within each quintile of the entire task, such that only the two last phases showed their significant association (bin 1: r = 0.081, *P* = 0.73; bin2: r = 0.22, *P* = 0.33; bin 3: r = 0.32, *P* = 0.16; bin 4, r = 0.57, *P* = 0.0074; bin 5, r = 0.43, *P* = 0.051; **Fig. 3d**). These results suggest that independent of individuals’ initial working memory capacity, how fast and to what extent individuals get trained at their working memory capacity may be associated with their behavioral preference for delayed rewards. See **figure S3** for the correlation results between performances at other intervention tasks and individuals’ impulsivity changes.

There was a stark difference in individuals’ behavioral pattern during the Control_WM_ task (i.e., spatial attention task). Across the task, individuals’ task performance (accuracy) gradually decreased (Pearson correlation, r = −0.36, *P* = 0.0011, **Fig. S4d**); 66.67% of participants showed decreasing task performance over time. Given this pattern, it is tempting to interpret these results as evidence that participants experienced mental fatigue over an easy and laborious task (Kluger, Krupp, & Enoka, 2013) than being trained for a specific type of executive function (e.g., selective attention). However, we cannot neglect an alternative possible explanation that participants’ task performance reflects a ceiling effect due to the low task difficulty, which introduces a potential bias toward finding a decremental effect. Nevertheless, we directly examined the impact of the task on individuals’ impulsivity, and found that neither mean task accuracy nor the performance slope (estimated speed of task performance change) of the Control_WM_ was correlated with the extent to which individuals’ delay discounting decrease (**Fig. S3d, S4e, S4f**). Together, there was no clear evidence suggesting that participating in a continuous attention task has an effect of reducing individuals’ impulsivity.

## 4. Experiment 2

### 4.1. Methods

Experiment 2 was a follow-up study that was designed to test whether or not the results from the first experiment could be replicated. Participants were randomly assigned to one of the three intervention tasks: working memory task and two control tasks. 30 students participated in the Working memory task. For control tasks, comparable number of students were assigned who were age, sex, and education matched (Control_WM_: N = 30; Resting: N = 30). 11 participants were excluded from the analysis because individual-level computational model did not arrive at a unique solution using MLE. In addition, nine participants who had discount rate estimates larger or smaller than 3 median absolute deviations (MAD) were excluded as outliers. After exclusion, data from 70 participants (male/female = 41/29, age = 22.17±4.08) were used for analyses (see **Table 2** for demographic information for each subgroup).

**Table 2.**
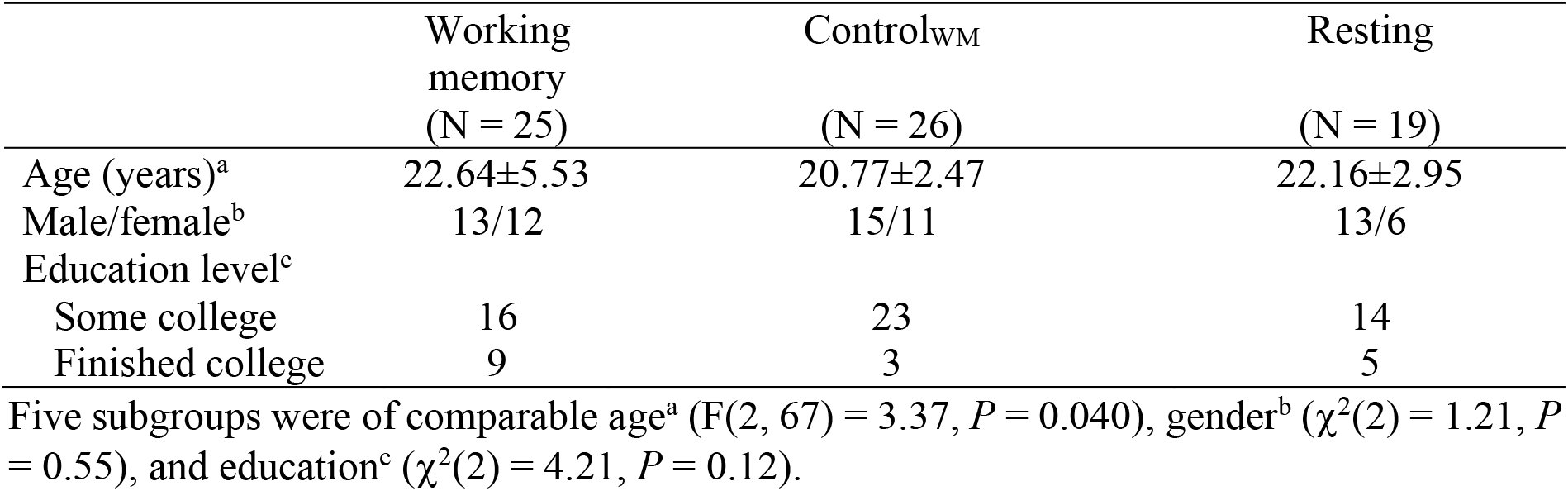
Demographic data for Experiment 2.

### 4.2. Results

In Experiment 1, we observed partial evidence suggesting that working memory enhancement may reduce individuals’ impulsivity. To further examine whether these results regarding the effect of working memory training on impulsivity reduction are replicable, we conducted an additional independent experiment where 70 participants (male/female = 41/29, age = 22.17±4.08; no overlap with Experiment 1) were asked to play either the Working memory or the Spatial attention tasks (Control_WM_), or to rest between two assessments of their impulsivity. As in Experiment 1, the three sub-groups showed comparable delay discount rates (F(2, 67) = 0.35, *P* = 0.71, one-way ANOVA; **Fig. 4**), which indicates that individuals’ baseline impulsivity levels were matched across groups.

**Figure 4.**
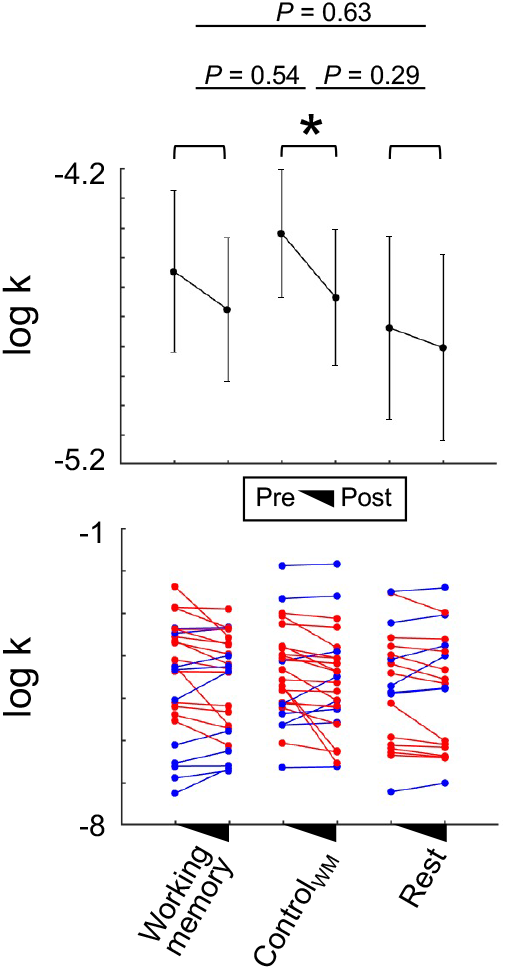
Impacts of the intervention tasks on reducing individuals’ impulsivity (Experiment 2). In Experiment 2, three types of intervention task (Working memory, Control_WM_, Rest) was tested. To examine whether each intervention had an impact on individuals’ impulsivity, we calculated changes in individuals’ delay discount rates (*k*) by comparing their behavioral patterns during the ITC task conducted before and after the intervention task. Individuals showed significantly reduced impulsivity after participating in the Control_WM_ (Paired t-test, t(20) = 2.07, *P* = 0.049, Cohen’s d = 0.41, bootstrapping *P* = 0.051). On the contrary, individuals who participated in other intervention tasks including Working memory task did not show the same pattern (all bootstrapping *p*s>0.05). The mean discount rate change in individuals who participated in the Control_WM_ task was not larger compared with that in individuals with any intervention task (all bootstrapping *p*s>0.05). Error bars indicate s.e.m.; *† < 0.10.

We analyzed participants’ choice data following the same protocol as used in Experiment 1. Individuals’ discount rate *k* were estimated from their behavioral choices as their impulsivity measure, and we used Δlog *k* as an indication of the impacts of intervention task. First, we tested the impacts of intervention tasks on individuals’ impulsivity reduction. Consistent with Experiment 1, we observed a significant main effect of Time (pre-versus post-intervention; F(1, 67) = 6.81, *P* = 0.011), but no significant effect of Type (Working memory, Control_WM_, and Rest; F(2, 67) = 0.22, *P* = 0.81) nor the interaction of Time × Type (F(2, 67) = 0.59, *P* = 0.56). Model-agnostic analyses on individuals’ choices were largely the same as these modelbased results (see **Fig. S5** for statistical results). Both experiments together, these results indicate that there is a significant repetition effect in reducing individuals’ impulsivity, while no evidence that supports impacts of intervention tasks.

Examining individuals’ impulsivity changes (changes in discount rates) between two assessments within each intervention group showed inconsistent patterns compared with Experiment 1. Specifically, the group who took part in the Working memory task or rested did not show significant effect of the intervention task (Working memory: Paired t-test, t(25) = 1.37, *P* = 0.18, Cohen’s d = 0.27, bootstrapping *P* = 0.18; Rest: t(19) = 0.86, *P* = 0.39, Cohen’s d = 0.20, bootstrapping *P* = 0.40), while the Control_WM_ group showed a significant reduction in their impulsivity (t(26) = 2.07, *P* = 0.047, Cohen’s d = 0.41, bootstrapping *P* = 0.048). Given these inconsistencies, one may suspect that our null results regarding the type of intervention task are due to lack of power. However, this is unlikely given that the results hold consistent even when we collapsed the data from both experiments (Time: F(1, 125) = 13.11, *P* = 4.25e-04; Type: F(2, 125) = 0.27, *P* = 0.77; Time × Type: F(2, 125) = 0.15, *P* = 0.86).

Second, we examined the association between individuals’ working memory task performances and their levels of impulsivity reduction. Note that consistent with Experiment 1, individuals showed performance increase along the course of the Working memory task (r = 0.50, *P* = 0.0017; **Fig. 5a**). In contrast to the association observed in the first experiment, neither the average accuracy in the Working memory task (r = −0.18, *P* = 0.38; **Fig. 5b**) nor the speed of performance improvement (r = −0.19, *P* = 0.35; **Fig. 5c**) showed significant correlation with the change of individuals’ discount rate. Moreover, gradual increase of the association between task improvement and impulsivity reduction, which we interpreted as evidence for the impact of the intervention task, disappeared in the replication data (**Fig. 5d**). The association between individuals’ performances in the Control_WM_ task and their impulsivity changes was also examined, albeit the results did not provide any clear evidence suggesting its impact on impulsivity reduction as reported in Experiment 1 (**Fig. S6b, S7e, S7f**).

**Figure 5.**
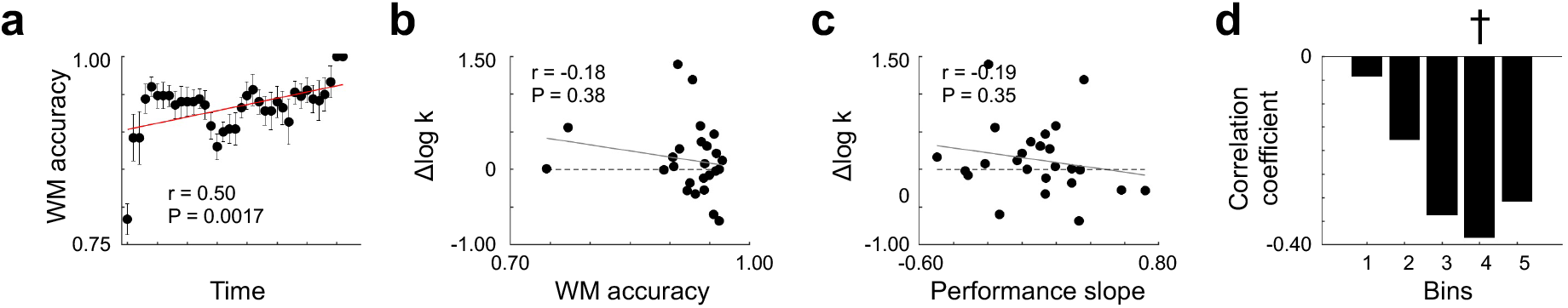
Individual performance enhancement in working memory training is not associated with the level of impulsivity reduction (Experiment 2). **(a)** Participants showed a significant performance improvement in Working memory task over time (Pearson’s correlation r = 0.50, *P* = 0.0017). Each dot represents mean accuracy within each window across participants, and error bars indicate s.e.m. **(b)** Individuals’ overall working memory performance was not associated with the changes in their delay discounting rates (r = −0.18, *P* = 0.38, bootstrapping *P* = 0.36). **(c)** Individuals’ speed of Working memory performance enhancement (performance slope) was not correlated with the level of impulsivity reduction (r = −0.19, *P* = 0.35, bootstrapping *P* = 0.35). Each dot represents an individual, and solid red lines are robust regression lines. **(d)** Over the course of the working memory task, no significant association was observed between individual’s Working memory accuracy and their impulsivity reduction level, and if any, there were negative trends which show that individuals tended to show more impulsive choices in the post-intervention ITC task. † < 0.10.

## 5. Discussion

The current study focused on two potential factors that are closely linked to successful future simulation and examined their impacts on modulating individuals’ impulsivity. Through two independent experiments, we found no consistent evidence supporting that either of cognitive intervention tasks has a specific and reliable impact of reducing individuals’ impulsivity. On the contrary to our original hypotheses, our results showed significant reduction of individuals’ impulsivity at the second assessment, regardless of the type of intervention task administered between two ITC tasks.

Mental simulation is suggested as one of the four modes of future thinking (Szpunar, Spreng, & Schacter, 2014), and broadly defined as cognitive construction of hypothetical scenarios or the reconstruction of real events. As a high-level cognitive function, future thinking is known to be crucial in planning, decision-making, and learning (Addis, Wong, & Schacter, 2007; Boyer, 2008; Buckner & Carroll, 2007; Na et al., 2021; Schacter & Addis, 2007; S. E. Snider, LaConte, & Bickel, 2016; Szpunar, Watson, & McDermott, 2007). Recent studies showed that such future thinking also plays an important role in evaluating temporally delayed rewards (Benoit et al., 2011; Peters & Büchel, 2010). Specifically, individuals preferred delayed rewards more than they used to after being enforced to focus on episodic future thoughts (Peters & Büchel, 2010). Given its multifaceted cognitive processes recruited through future thinking, it was premature to conclude why individuals’ behavioral preferences changed under manipulation. We examined two potential mechanisms which may underlie or related to episodic future thinking to further our understanding regarding individual differences along their preference over delayed rewards. By training and enhancing cognitive abilities related to each mechanism, we expected individuals’ mental simulation to become more fluent and in turn, contribute to impulsivity reduction. In contrast to our expectation, across two independent tasks, we did not find any specific modulation effects from either of targeted cognitive intervention tasks. These results may be due to complex nature of future oriented thinking process which cannot be separated into independent components. Thus, each type of intervention tested in the current study, without being paired up with other complementary tasks, did not have either strong or reliable modulatory effects.

Impacts of working memory training for reducing individuals’ impulsivity have been inconclusive. For example, delay discounting tendency of individuals who have stimulant use disorder was significantly reduced after multiple sessions of working memory training (Bickel et al., 2011). In other sample, however, working memory training was effective only for the condition where participants were explicitly cued to execute episodic future thinking (Sarah E Snider et al., 2018). We speculated that such an inconsistency may be related to subtle differences in working memory task designs; only the former study required numerical processing (i.e., digit span). However, our two independent experiments which both required numerical working memory from individuals showed mixed results. Given the short training period that our study implemented, we do not claim that *all* working memory trainings are ineffective in impulsivity reduction, but note that previously suggested intervention paradigms should also be revisited.

Taking two independent experimental data together, individuals’ reduced impulsivity observed in the current study was not linked to a specific type of intervention task, but rather appeared as general phenomena. These results suggest that our study design may be triggering cognitive and affective mechanisms over the repeated applications of ITC task. Such malleability of intertemporal choice has been documented across various contexts (Lempert & Phelps, 2016). For example, attending to a particular type of information (e.g., magnitude of rewards) or updating one’s expectation based on recent experiences are known to affect individuals’ impulsivity (Ariely & Loewenstein, 2000; Ebert & Prelec, 2007; Loewenstein, 1988). In the current study, participants already went through entire series of choices between delayed and immediate options before intervention tasks, which could have affected individuals’ expectation about the reward delays. Participants would perceive reward delays less distant when expecting a non-zero days of delay as a new ‘reference’ delay (Loewenstein, 1988) at the beginning of the second assessment, and in turn, show less impulsive choices. We unfortunately did not collect a direct measure of individuals’ expected delays, and thus our suggested interpretation needs further investigation.

There are a few alternative explanations why individuals showed reduced impulsivity at the second assessment independent of the type of intervention task. First, because individuals already tried simulating future rewards once, simulation of the value of delayed rewards might had been facilitated. Previously, it was shown that individuals who could vividly project one’s previous experience into the future showed particularly large impulsivity reduction from episodic future thinking (Peters & Büchel, 2010). Our intervention tasks between two ITC tasks lasted around 20 minutes, and we cannot fully rule out the possibility that individuals remembered their first choices and viewed options, albeit the choice sequences were randomized. Note that this possibility should be considered apart from a simple practice effect, because there was no evidence of impulsivity reduction along three repeated blocks within each experiment (Exp1: F(2, 581) = 1.78, *P* = 0.17; Exp2: F(2, 415) = 0.52, *P* = 0.60). Second, participants might had felt bored or stressed out at the later stage of the experiment; besides cognitive context manipulations, being under specific emotional states is also known to make individuals more or less impulsive (DeSteno, Li, Dickens, & Lerner, 2014; Guan, Cheng, Fan, & Li, 2015). Third, letting individuals to spend a period of time participating in other types of cognitive tasks (or mind wondering) might had led them to think about thoughts unrelated to delayed rewards and had an impact of reducing delay discounting (Smallwood et al., 2011; Smallwood, Ruby, & Singer, 2013). Future studies can be tailored to directly test each hypothesis and disambiguate the mechanisms explaining why individuals’ impulsivity decrease at a repeated assessment.

The current study has some limitation that needs further investigation. First, null results from the second experiment might be due to small sample size and lack statistical power. This is unlikely to be a main factor because the results regarding the impact of working memory training did not hold when both data set from two experiments were pooled together. Second, we only examined the impacts of fixed doses of intervention tasks, and thus we cannot completely rule out that a larger dose (longer training and repetition) would induce significant impulsivity reduction effects. Third, in contrast to the first experiment, the second experiment was conducted during the COVID-19 pandemic (Exp1: March 25, 2019-May 20, 2019; Exp2: January 13, 2021-July 19, 2021), and individuals in general are expected to be in negative mental states (e.g., high stress, depression, and anxiety) (Brooks et al., 2020; Pfefferbaum & North, 2020). Given the close relationship between the on-going pandemic and health risk behaviors (Clay & Parker, 2020; Johnson et al., 2022; Park, Lee, Sul, & Chung, 2021), we cannot overlook a possibility that the null effect of intervention training at the second experiment is associated with the environmental change and accompanied changes of mental states.

Nevertheless, the current study delineated two possible mechanisms explaining the positive relationship between episodic future thinking and changes in delay discounting behavior. Based on two independent experiments, no specific task among the examined intervention tasks showed a reliable impact of reducing individuals’ impulsivity. Rather unexpectedly, we observed a type-general impulsivity reduction effect. The current work leaves it an open question how to design cognitive training paradigms that target more specific cognitive functions for effective and efficient intervention to individuals who have health risk problems, such as obesity (Davis, Patte, Curtis, & Reid, 2010; Jarmolowicz et al., 2014; Stoeckel, Murdaugh, Cox, Cook, & Weller, 2013), substance addiction (Han et al., 2015; Kirby & Petry, 2004; MacKillop et al., 2011), and pathological gambling (Dixon et al., 2003; Power, Goodyear, & Crockford, 2012). Still, these results once more point out that an individual’s tendency to act impulsively may not be a trait-like attribute that does not change, but a malleable feature that can be changed differently depending on the situated contexts (e.g., education, and socioeconomic status of the family (Watts, Duncan, & Quan, 2018)).

## Acknowledgments

We gratefully acknowledge assistance from the Decision Neuroscience and Cognitive Engineering lab members. This work was supported in part by the National Research Foundation of Korea (NRF-2018R1D1A1B07043582, NRF-2018M3C1B8013691 to Chung).

## Author contributions

M.H., S.-P.K., and D.C. designed the experiments. M.H. and D.C. analyzed the data. M.H. and D.C. drafted the manuscript. All of the authors discussed the results, revised, and approved the final manuscript.

## Data availability

Analytic scripts and data are available on GitHub (https://github.com/dongilchung/impulsivity-intervention).

## References

Addis, D. R., Wong, A. T., & Schacter, D. L. (2007). Remembering the past and imagining the future: common and distinct neural substrates during event construction and elaboration. Neuropsychologia, 45(7), 1363–1377.

Ainslie, G. (1975). Specious reward: a behavioral theory of impulsiveness and impulse control. Psychological bulletin, 82(4), 463.

Ariely, D., & Loewenstein, G. (2000). When does duration matter in judgment and decision making? Journal of Experimental Psychology: General, 129(4), 508.

Azfar, O. (1999). Rationalizing hyperbolic discounting. Journal of Economic Behavior & Organization, 38(2), 245–252.

Baker, F., Johnson, M. W., & Bickel, W. K. (2003). Delay discounting in current and never-before cigarette smokers: similarities and differences across commodity, sign, and magnitude. Journal of abnormal psychology, 112(3), 382.

Benoit, R. G., Gilbert, S. J., & Burgess, P. W. (2011). A neural mechanism mediating the impact of episodic prospection on farsighted decisions. Journal of Neuroscience, 31(18), 6771–6779.

Bickel, W. K., & Marsch, L. A. (2001). Toward a behavioral economic understanding of drug dependence: delay discounting processes. Addiction, 96(1), 73–86.

Bickel, W. K., Wilson, A. G., Franck, C. T., Mueller, E. T., Jarmolowicz, D. P., Koffarnus, M. N., & Fede, S. J. (2014). Using crowdsourcing to compare temporal, social temporal, and probability discounting among obese and non-obese individuals. Appetite, 75, 82–89.

Bickel, W. K., Yi, R., Landes, R. D., Hill, P. F., & Baxter, C. (2011). Remember the future: working memory training decreases delay discounting among stimulant addicts. Biological psychiatry, 69(3), 260–265.

Boyer, P. (2008). Evolutionary economics of mental time travel? Trends Cogn Sci, 12(6), 219–224. doi:10.1016/j.tics.2008.03.003

Brooks, S. K., Webster, R. K., Smith, L. E., Woodland, L., Wessely, S., Greenberg, N., & Rubin, G. J. (2020). The psychological impact of quarantine and how to reduce it: rapid review of the evidence. The lancet, 395(10227), 912–920.

Brown, S. W. (1997). Attentional resources in timing: Interference effects in concurrent temporal and nontemporal working memory tasks. Perception & psychophysics, 59(7), 1118–1140.

Buckner, R. L., & Carroll, D. C. (2007). Self-projection and the brain. Trends in cognitive sciences, 11(2), 49–57.

Clay, J. M., & Parker, M. O. (2020). Alcohol use and misuse during the COVID-19 pandemic: a potential public health crisis? The Lancet. Public Health, 5(5), e259.

D’Argembeau, A., Ortoleva, C., Jumentier, S., & Van der Linden, M. (2010). Component processes underlying future thinking. Memory & cognition, 38(6), 809–819.

Davis, C., Patte, K., Curtis, C., & Reid, C. (2010). Immediate pleasures and future consequences. A neuropsychological study of binge eating and obesity. Appetite, 54(1), 208–213. doi:10.1016/j.appet.2009.11.002

DeSteno, D., Li, Y., Dickens, L., & Lerner, J. S. (2014). Gratitude: A tool for reducing economic impatience. Psychological science, 25(6), 1262–1267.

Dixon, M. R., Marley, J., & Jacobs, E. A. (2003). Delay discounting by pathological gamblers. Journal of applied behavior analysis, 36(4), 449–458.

Ebert, J. E., & Prelec, D. (2007). The fragility of time: Time-insensitivity and valuation of the near and far future. Management science, 53(9), 1423–1438.

Figner, B., Knoch, D., Johnson, E. J., Krosch, A. R., Lisanby, S. H., Fehr, E., & Weber, E. U. (2010). Lateral prefrontal cortex and self-control in intertemporal choice. Nat Neurosci, 13(5), 538–539. doi:10.1038/nn.2516

Frederick, S., Loewenstein, G., & O’donoghue, T. (2002). Time discounting and time preference: A critical review. Journal of economic literature, 40(2), 351–401.

Green, L., & Myerson, J. (2004). A discounting framework for choice with delayed and probabilistic rewards. Psychol Bull, 130(5), 769–792. doi:10.1037/0033-2909.130.5.769

Guan, S., Cheng, L., Fan, Y., & Li, X. (2015). Myopic decisions under negative emotions correlate with altered time perception. Frontiers in psychology, 6, 468.

Hakimi, S., & Hare, T. A. (2015). Enhanced neural responses to imagined primary rewards predict reduced monetary temporal discounting. Journal of Neuroscience, 35(38), 13103–13109.

Han, R., Takahashi, T., Miyazaki, A., Kadoya, T., Kato, S., & Yokosawa, K. (2015). Activity in the left auditory cortex is associated with individual impulsivity in time discounting. Paper presented at the 2015 37th Annual International Conference of the IEEE Engineering in Medicine and Biology Society (EMBC).

Hare, T. A., Camerer, C. F., & Rangel, A. (2009). Self-control in decision-making involves modulation of the vmPFC valuation system. Science, 324(5927), 646–648.

Hayman, M. (1942). Two minute clinical test for measurement of intellectual impairment in psychiatric disorders. Archives of Neurology & Psychiatry, 47(3), 454–464.

Hill, P. F., & Emery, L. J. (2013). Episodic future thought: Contributions from working memory. Consciousness and cognition, 22(3), 677–683.

Jarmolowicz, D. P., Cherry, J. B., Reed, D. D., Bruce, J. M., Crespi, J. M., Lusk, J. L., & Bruce, A. S. (2014). Robust relation between temporal discounting rates and body mass. Appetite, 78, 63–67. doi:10.1016/j.appet.2014.02.013

Johnson, S. L., Porter, P., Modavi, K., Dev, A., Pearlstein, J., & Timpano, K. R. (2022). Emotion-related impulsivity predicts increased anxiety and depression during the COVID-19 pandemic. Journal of affective disorders.

Kable, J. W., & Glimcher, P. W. (2007a). The neural correlates of subjective value during intertemporal choice. Nature neuroscience, 10(12), 1625.

Kable, J. W., & Glimcher, P. W. (2007b). The neural correlates of subjective value during intertemporal choice. Nature neuroscience, 10(12), 1625–1633.

Kirby, K. N., & Petry, N. M. (2004). Heroin and cocaine abusers have higher discount rates for delayed rewards than alcoholics or non-drug-using controls. Addiction, 99(4), 461–471.

Klingberg, T. (2010). Training and plasticity of working memory. Trends in cognitive sciences, 14(7), 317–324.

Kluger, B. M., Krupp, L. B., & Enoka, R. M. (2013). Fatigue and fatigability in neurologic illnesses: proposal for a unified taxonomy. Neurology, 80(4), 409–416.

Laibson, D. (1997). Golden eggs and hyperbolic discounting. The Quarterly Journal of Economics, 112(2), 443–478.

Lempert, K. M., & Phelps, E. A. (2016). The malleability of intertemporal choice. Trends in cognitive sciences, 20(1), 64–74.

Loewenstein, G. F. (1988). Frames of mind in intertemporal choice. Management science, 34(2), 200–214.

MacKillop, J., Amlung, M. T., Few, L. R., Ray, L. A., Sweet, L. H., & Munafo, M. R. (2011). Delayed reward discounting and addictive behavior: a meta-analysis. Psychopharmacology (Berl), 216(3), 305–321. doi:10.1007/s00213-011-2229-0

Mazur, J. E. (1987). An adjusting procedure for studying delayed reinforcement. Commons, ML.; Mazur, JE.; Nevin, JA, 55–73.

McClure, S. M., Laibson, D. I., Loewenstein, G., & Cohen, J. D. (2004). Separate neural systems value immediate and delayed monetary rewards. Science, 306(5695), 503–507.

Na, S., Chung, D., Hula, A., Perl, O., Jung, J., Heflin, M., … Gu, X. (2021). Humans use forward thinking to exploit social controllability. eLife, 10.

Park, J., Lee, S., Sul, S., & Chung, D. (2021). Depression symptoms mediate mismatch between perceived severity of the COVID-19 pandemic and preventive motives. Frontiers in psychology, 12.

Peters, J., & Büchel, C. (2010). Episodic future thinking reduces reward delay discounting through an enhancement of prefrontal-mediotemporal interactions. Neuron, 66(1), 138–148.

Petry, N. M., & Casarella, T. (1999). Excessive discounting of delayed rewards in substance abusers with gambling problems. Drug and alcohol dependence, 56(1), 25–32.

Pfefferbaum, B., & North, C. S. (2020). Mental health and the Covid-19 pandemic. New England Journal of Medicine, 383(6), 510–512.

Power, Y., Goodyear, B., & Crockford, D. (2012). Neural correlates of pathological gamblers preference for immediate rewards during the iowa gambling task: an fMRI study. J Gambl Stud, 28(4), 623–636. doi:10.1007/s10899-011-9278-5

Schacter, D. L., & Addis, D. R. (2007). The cognitive neuroscience of constructive memory: remembering the past and imagining the future. Philosophical Transactions of the Royal Society B: Biological Sciences, 362(1481), 773–786.

Schacter, D. L., Addis, D. R., Hassabis, D., Martin, V. C., Spreng, R. N., & Szpunar, K. K. (2012). The future of memory: remembering, imagining, and the brain. Neuron, 76(4), 677–694.

Smallwood, J., Brown, K. S., Tipper, C., Giesbrecht, B., Franklin, M. S., Mrazek, M. D., … Schooler, J. W. (2011). Pupillometric evidence for the decoupling of attention from perceptual input during offline thought. PloS one, 6(3), e18298.

Smallwood, J., Ruby, F. J., & Singer, T. (2013). Letting go of the present: mind-wandering is associated with reduced delay discounting. Consciousness and cognition, 22(1), 1–7.

Snider, S. E., Deshpande, H. U., Lisinski, J. M., Koffarnus, M. N., LaConte, S. M., & Bickel, W. K. (2018). Working memory training improves alcohol users’ episodic future thinking: A rate-dependent analysis. Biological Psychiatry: Cognitive Neuroscience and Neuroimaging, 3(2), 160–167.

Snider, S. E., LaConte, S. M., & Bickel, W. K. (2016). Episodic Future Thinking: Expansion of the Temporal Window in Individuals with Alcohol Dependence. Alcohol Clin Exp Res, 40(7), 1558–1566. doi:10.1111/acer.13112

Stoeckel, L. E., Murdaugh, D. L., Cox, J. E., Cook, E. W., 3rd, & Weller, R. E. (2013). Greater impulsivity is associated with decreased brain activation in obese women during a delay discounting task. Brain Imaging Behav, 7(2), 116–128. doi:10.1007/s11682-012-9201-4

Story, G., Vlaev, I., Seymour, B., Darzi, A., & Dolan, R. (2014). Does temporal discounting explain unhealthy behavior? A systematic review and reinforcement learning perspective. Frontiers in behavioral neuroscience, 8, 76.

Szpunar, K. K. (2010). Episodic future thought: An emerging concept. Perspectives on Psychological Science, 5(2), 142–162.

Szpunar, K. K., Spreng, R. N., & Schacter, D. L. (2014). A taxonomy of prospection: Introducing an organizational framework for future-oriented cognition. Proceedings of the National Academy of Sciences, 111(52), 18414–18421.

Szpunar, K. K., Watson, J. M., & McDermott, K. B. (2007). Neural substrates of envisioning the future. Proceedings of the National Academy of Sciences, 104(2), 642–647.

Watts, T. W., Duncan, G. J., & Quan, H. (2018). Revisiting the marshmallow test: A conceptual replication investigating links between early delay of gratification and later outcomes. Psychological science, 29(7), 1159–1177.

Zavagnin, M., De Beni, R., Borella, E., & Carretti, B. (2016). Episodic future thinking: the role of working memory and inhibition on age-related differences. Aging clinical and experimental research, 28(1), 109–119.

