## Supplementary material for "Simulating future rewards: exploring the impacts of implicit context association and arithmetic booster in delay discounting"

### for

### Supplementary Figures and Legends

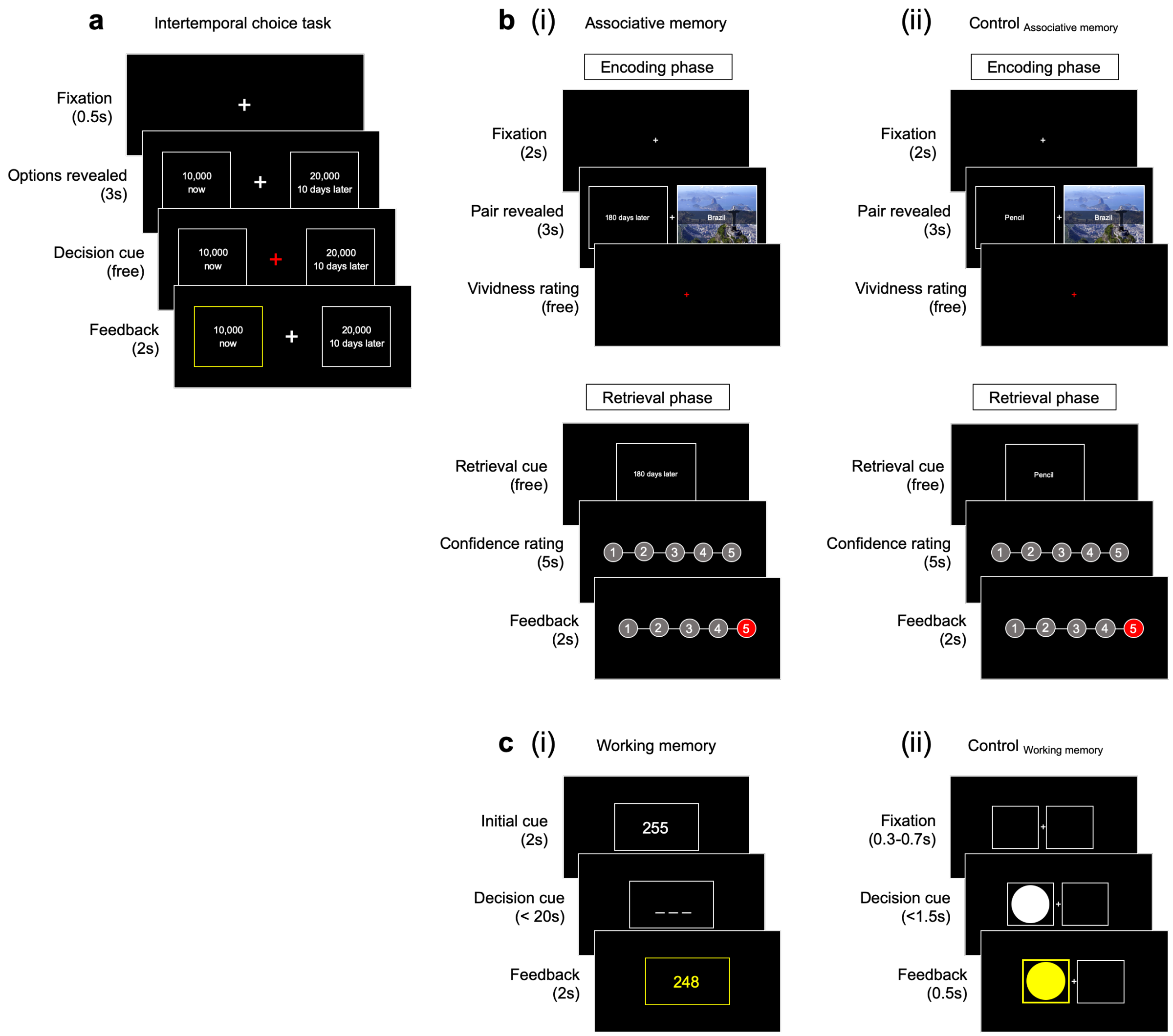


**Figure S1. Task timeline for each paradigm. (a)** Each trial started with a crosshair screen. Participants made a series of choices between two options; one option was always an immediate small reward whereas the other option was a delayed large reward. A red crosshair on the center of the screen was used as a decision cue that informed participants to make a choice. The selected option was highlighted with a yellow bordered box at the feedback presentation. **(bi, bii)** Associative memory task and its matched control task (Control$_{\text{Associative memory}}$) consisted of 'encoding' and 'retrieval' phases. During the encoding phase, a set of pairs between temporal delays (or stationaries for the Control$_{\text{Associative memory}}$ task) and names of the countries were presented. Each name of the country was presented on top of a picture that showed one of must-see sights within the country. A red crosshair on the center of the screen was used as a cue asking participants to report whether or not they could vividly imagine the paired association. During the retrieval phase, participants were presented with cues (temporal delays for Associative memory and stationaries for Control$_{\text{AM}}$) and asked to verbally report the name of the country that was paired during the encoding phase. Furthermore, participants were asked to report how confident they were of the submitted answer before moving on to the next retrieval cue. **(ci, cii)** Information regarding the Working memory task and Control$_{\text{working memory}}$ task is identical to that in **figure 1**.

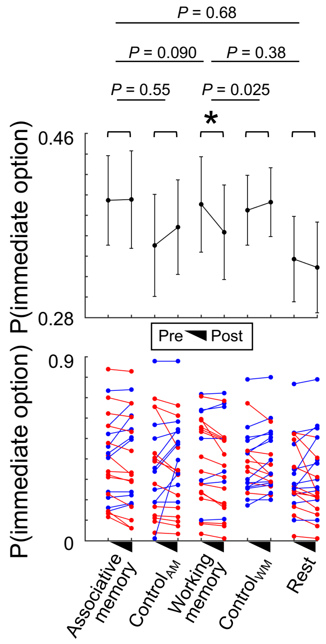


**Figure S2. Impacts of the intervention tasks on reducing individuals' impulsivity (Experiment 1).** For completeness, we also compared the impacts of each intervention design using model-agnostic measures; ΔP(immediate option) = P(immediate option$_{\text{pre-intervention}}$) − P(immediate option$_{\text{post-intervention}}$) was calculated as a model-agnostic measure where P(immediate option) refers to probability of choosing the immediate option. Among the three intervention groups (Associative memory, Working memory, and Rest), no main or interaction effects of Time (within-group) and Type (between-group) factors were found significant in model-agnostic measures (Mixed-design ANOVA; Time: F(1, 64) = 1.89, *P* = 0.17; Type: F(2, 64) = 0.65, *P* = 0.52; Interaction: F(2, 64) = 1.02, *P* = 0.37). Within each intervention group, the Working memory task was the only intervention that showed a significant reduction in individuals' impulsive choices (Paired t-test, t(20) = 2.62, *P* = 0.017, Cohen's d = 0.57, bootstrapping *P* = 0.016); other interventions besides the Working memory task all failed to show significant change in participants' choice behaviors (Associative memory: t(20) = -0.058, *P* = 0.95, Cohen's d = -0.012, bootstrapping *P* = 0.96; Control$_{AM}$: t(21) = -0.73, *P* = 0.48, Cohen's d = -0.15, bootstrapping *P* = 0.47; Control$_{WM}$: t(20) = -0.70, *P* = 0.49, Cohen's d = -0.15, bootstrapping *P* = 0.49; Rest: t(22) = 0.46, *P* = 0.65, Cohen's d = 0.10, bootstrapping *P* = 0.66). Although impacts of the intervention types were statistically comparable, the Working memory task had a relatively larger effect in reducing participants' tendency of choosing the immediate option than the Associative memory task (Two-sample t-test, t(42) = 1.71, *P* = 0.095, Cohen's d = 0.26, bootstrapping *P* = 0.090) or the Rest (t(42) = 0.91, *P* = 0.37, Cohen's d = 0.14, bootstrapping *P* = 0.38). Compared with a matched control intervention task (Control$_{WM}$), the Working memory intervention had a superior impact on individuals' choice tendency (t(40) = 2.30, *P* = 0.027, Cohen's d = 0.35, bootstrapping *P* = 0.025). Furthermore, consistent with model-based results, the Associative memory intervention showed comparable impacts compared with Control$_{AM}$ (t(43) = 0.63, *P* = 0.53, Cohen's d = 0.094, bootstrapping *P* = 0.55) or the Rest (t(44) = -0.41, *P* = 0.68, Cohen's d = -0.061, bootstrapping *P* = 0.68). Error bars indicate s.e.m.; $^{*}P < 0.05$.

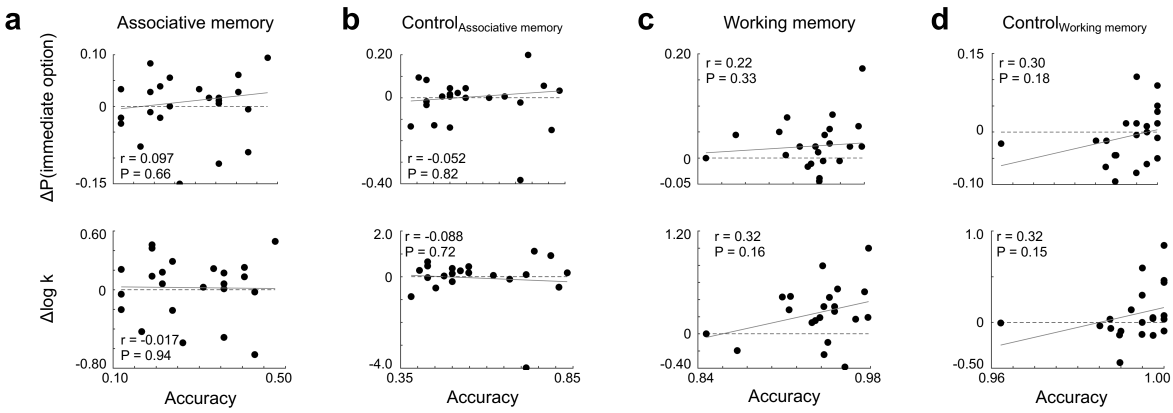


**Figure S3. Correlation between performance for each intervention task and individuals' impulsivity change (Experiment 1).** For completeness, correlations were examined between task performance and individuals' impulsivity change for all four types of intervention tasks. **(a, b)** Neither the Associative memory nor Control$_{AM}$ showed significant relationship between task performance and individuals' impulsivity changes. **(c)** Individuals who were assigned to the Working memory intervention task showed a trend of impulsivity reduction (positive $\Delta$P(immediate option) or $\Delta$log k; top: model-agnostic, Pearson's correlation, r = 0.22, $P = 0.34$, bootstrapping $P = 0.33$; bottom: model-based, r = 0.32, $P = 0.16$, bootstrapping $P = 0.16$; copy of **Fig. 3c**). **(d)** The intervention task performance from individuals who participated in the Control$_{WM}$ task also showed non-significant relationship with the extent to which their impulsivity reduced (top : model-agnostic, r = 0.30, $P = 0.18$, bootstrapping $P = 0.18$; bottom: model-free, r = 0.32, $P = 0.16$, bootstrapping $P = 0.15$). Each dot represents individual participant.

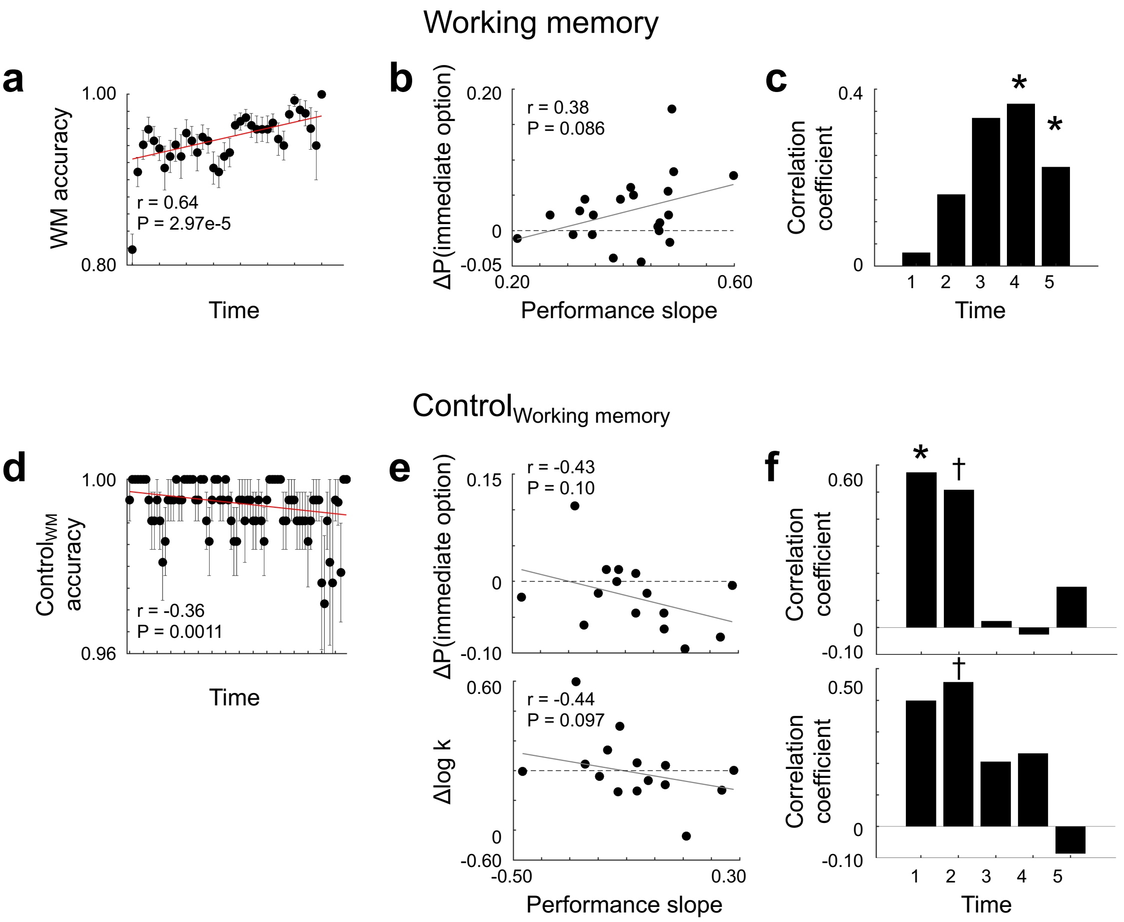


**Figure S4. Correlation between performance change during intervention tasks and individuals' impulsivity change (Experiment 1). (a)** Individuals showed a significant working memory performance improvement over time (Pearson's correlation, r = 0.64, *P* = 2.97e-5). **(b)** Consistent with model-based results (**Fig. 3c, 3d**), individuals who showed steeper performance enhancement in the Working memory task tend to show relatively larger impulsivity reduction (r = 0.38, *P* = 0.092, bootstrapping *P* = 0.086), and **(c)** this relationship between tasks was relatively more pronounced when the average performance at the later part of the intervention task was used for the correlation analyses. **(d)** Individuals' performance change in the Control$_{WM}$ was examined. Unlike the Working memory task, participants' task accuracy significantly decreased over time (r = -0.36, *P* = 0.0011). **(e)** There were negatively trending relationship between the steepness of performance change and individuals' impulsivity reduction in both model-agnostic (top) and model-based measures (bottom). **(f)** Mirroring the analysis methods for the Working memory task, the Control$_{WM}$ task was divided into five bins (bin size = 1/3 of the entire task length; sliding window with 1/6 overlap), and average task performance was calculated per each bin. Individuals' task performance at the early stage of the manipulation task was significantly correlated with the extent to which their impulsivity was reduced at the second ITC task. However, such a pattern disappeared for the task performance at the later stage (top: model-agnostic, bin 1 r = 0.57, *P* = 0.026; bin 2 r = 0.51, *P* = 0.053; bin 3 r = 0.023, *P* = 0.93; bin 4 r = -0.026, *P* = 0.93; bin 5 r = 0.15, *P* = 0.59; bottom: model-based, bin 1 r = 0.40, *P* = 0.14; bin 2 r = 0.46, *P* = 0.087; bin 3 r = 0.21, *P* = 0.46; bin 4 r = 0.23, *P* = 0.41; bin 5 r = -0.085, *P* = 0.76). $^{*}P < 0.05$. $^{*}$† < 0.10.

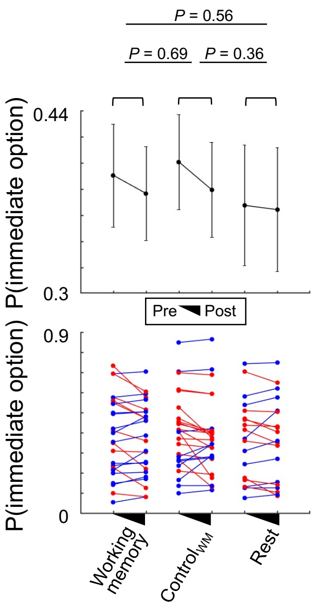


**Figure S5. Impacts of the intervention tasks on reducing individuals' impulsivity (Experiment 2).** For completeness, we also compared the impacts of each intervention design using model-agnostic measures. To examine whether each intervention had an impact on individuals' impulsivity, we calculated changes in their tendency of choosing the immediate option by comparing their behavioral patterns during the intertemporal choice (ITC) task conducted before and after each intervention task. Individuals did not show any significant changes of impulsivity after participating in the all intervention task (all bootstrapping *p*s>0.05). Also, individuals who participated in the Working memory task did not show significant impulsivity reduction compared with individuals who were assigned to other intervention task (all bootstrapping *p*s>0.05). Error bars indicate s.e.m.; $^{*}P < 0.05$.

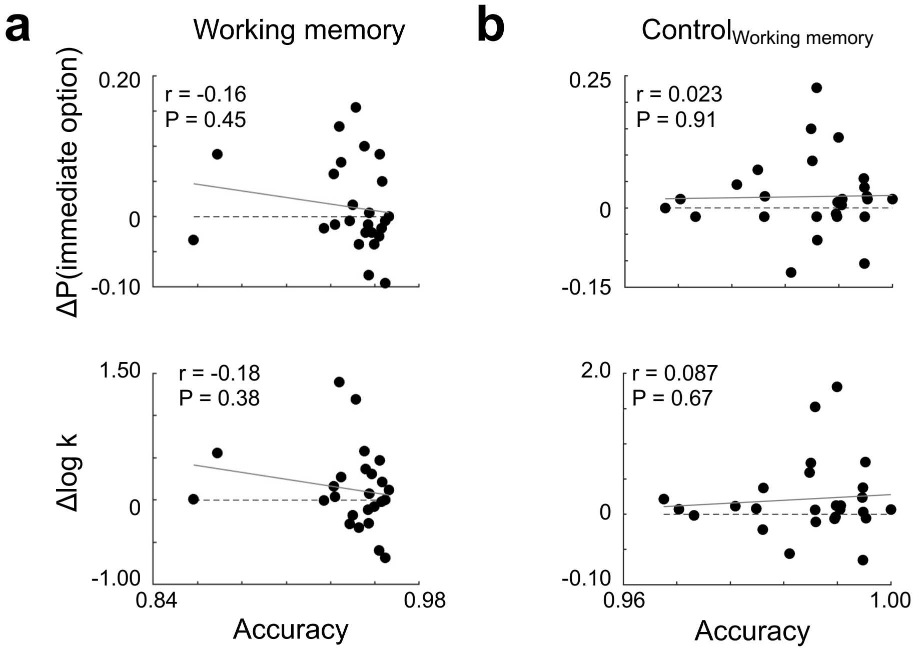


**Figure S6. Correlation between performance for each intervention task and individuals' impulsivity change (Experiment 2).** For completeness, correlations were examined between task performance and individuals' impulsivity change for both types of intervention tasks in Experiment 2. **(a, b)** Neither the Working memory (copy of **Fig. 5b**) nor Control$_{WM}$ showed significant relationship between task performance and individuals' impulsivity changes. Each dot represents individual participant.

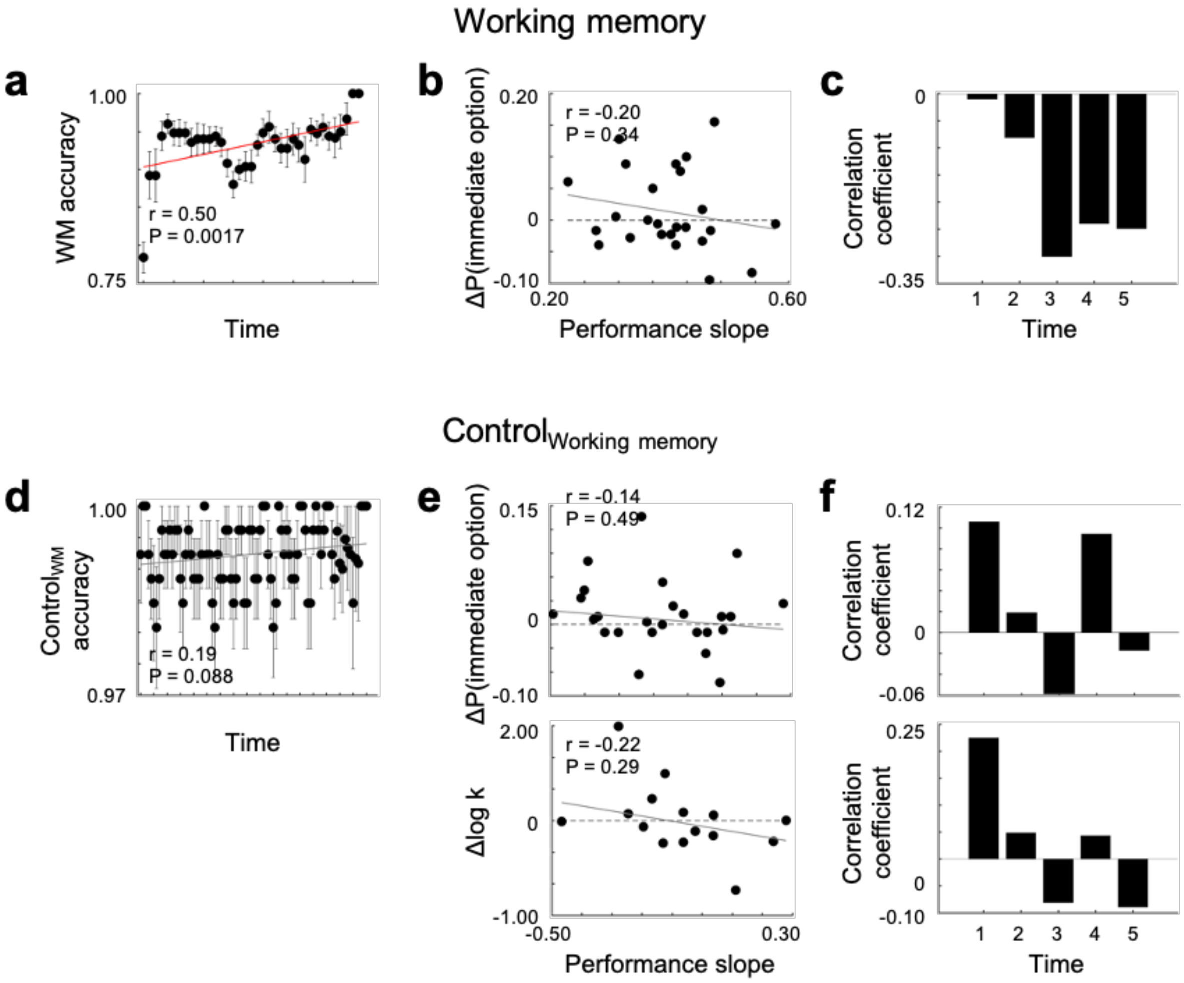


**Figure S7. Correlation between performance change during intervention tasks and individuals' impulsivity change (Experiment 2). (a)** Individuals showed a significant working memory performance improvement over time (Pearson's correlation, r = 0.19, *P* = 0.088). **(b, c)** Individuals' performance enhancement in the Working memory task did not show significant association with impulsivity reduction (r = -0.20, *P* = 0.34, bootstrapping *P* = 0.34). **(d)** Unlike the Working memory task, participants' task accuracy in the Control$_{WM}$ did not significantly change over time (r = 0.19, *P* = 0.088). **(e, f)** Individuals' task performance change was not correlated with the extent to which their impulsivity was reduced at the second ITC task.
